# Lymphatic vessel development in human embryos

**DOI:** 10.1101/2023.08.12.553102

**Authors:** Shoichiro Yamaguchi, Natsuki Minamide, Hiroshi Imai, Tomoaki Ikeda, Masatoshi Watanabe, Kyoko Imanaka-Yoshida, Kazuaki Maruyama

## Abstract

Lymphatic vessel development has been a subject of research for about 120 years. Studies employing mice and zebrafish models have elucidated that lymphatic endothelial cells (LECs) predominantly differentiate from venous endothelial cells via the expression of transcription factor Prospero homeobox protein 1 (Prox1), a master regulator of lymphatic vessel development. On the other hand, it has been found that LECs can also be generated from undifferentiated mesodermal or hemogenic endothelial cells, suggesting potential diversity in their origins depending on the organ or anatomical location. However, knowledge of human lymphatic vessel development remains limited. Here, we examined early lymphatic development in humans by analyzing 31 embryos and three 9-week old fetuses. We found that human embryos produce Prox1-expressing LECs in and around the cardinal veins, which converged to form initial lymph sacs. Furthermore, we also examined lymphatic vessel development in the heart, lungs, lower jaw, mesentery, intestines and kidneys. Lymphatic vessels appeared to develop at different rates in each organ and to display temporal differences in marker expression. These observation showed the possibility that there could exist different patterns of lymphatic vessel development across organs, which may reflect different cellular origins or developmental signaling in each organ.

Our research clarifies the early development of human lymphatic vessels, contributing to a better understanding of the evolution and phylogenetic relationships of lymphatic systems, and enriching our knowledge of the role of lymphatics in various human diseases.

**Significance Statement:** Lymphatic vessel development has been a focus of research for over a century. Recent studies across a variety of species have demonstrated that lymphatic endothelial cells originate from embryonic veins, and undifferentiated mesodermal cells. However, whether these findings are applicable to human has yet to be determined. In this study, we explored lymphatic vessel development in humans. Our analysis demonstrated that lymphatic endothelial cells in human embryos initially derived from embryonic veins. Notably, we found that lymphatic vessels in different organs displayed distinct developmental and marker expression patterns, suggesting a diversity in lymphatic vessel development across organs. Our research revealed the human lymphatic vessel development, contributing to the understanding of phylogenetics of lymphatic vessels and lymph-related diseases.

## Introduction

The lymphatic vascular system is essential for fluid transportation, immune reaction and lipids absorption. Lymphatic vessels are also important for various pathophysiological conditions(1– 3), including myocardial infarction(4–7), encephalitis(8), and Alzheimer’s disease(9). The development of lymphatic vessels has long been a subject of debate. Early studies conducted in the 1900s proposed that initial lymphatic vessels (lymph sacs) originate from venous endothelial cells and the entire lymphatic system subsequently proliferates from these structures into adjacent tissues and organs(10). An alternative theory suggested that lymph sacs originate from undifferentiated mesodermal cells and subsequently establish connections between jugular veins(11). An anatomical study on lymphatic development using human embryos was conducted by Putte in 1975(12). However, at that time, lymphatic markers had not been identified yet. As a result, it did not provide comprehensive detailed observations.

In recent years, progress in genetic analysis and the development of specific markers has significantly enhanced our understanding. The transcription factor Prospero homeobox protein 1 (Prox1)(13), a master regulator of lymphatic endothelial cell (LEC) differentiation, and the Vascular Endothelial Growth Factor C (VEGF-C)-VEGF Receptor 3 (VEGFR3) axis(14, 15), which is crucial for LECs proliferation, are essential molecules of the lymphatic vessel development. Genetic lineage analysis using mouse embryos with *Prox1-CreERT2* has clarified that LECs primarily originate from the cardinal veins(16). Furthermore, we have recently elucidated that in mice, LECs not only derived from venous endothelial cells, but in the head and neck region and the mediastinum, they may originate from evolutionarily conserved cardiopharyngeal mesoderm, which also generates musculatures and connective tissues in these regions(17). It has also been reported that the progenitors of lymphatic vessels may differ among organs, such as the heart(6, 18, 19), mesentery(20), and skin(21, 22).

Lymphatic malformations are one of the most common vascular disorders that occur in approximately one in 2,000-4,000 births(23). Lymphatic malformations tend to occur from the head and neck region to the mediastinum and are considered to be caused by abnormalities in the developmental process. Although mouse models of lymphatic malformations have been reported, it has been challenging to reproduce the anatomical characteristics of humans in mice(24). This might be because the lymphatic development in humans differs from that in the mice, but the exact cause remains poorly understood. Furthermore, each species possesses unique function and anatomy of the lymphatic system. For example, in tuna, the lymphatic vessels play a crucial role in moving the fins(25). In amphibians and reptiles, there are lymphatic hearts, which are part of lymphatic vessels that contract autonomously(26). While in birds, lymphatic hearts are once formed in embryonic periods, they disappear as the development progresses(27). Thus, the development, function, and anatomy of the lymphatic vessels display variations between species, and the knowledge obtained from mice does not necessarily apply to humans. Therefore, as a fundamental basis for studying human lymphatic diseases, it is crucial to understand lymphatic vessel development in humans.

In this study, we attempted to examine lymphatic vessel development in humans using 31 embryos and 3 fetuses at the 9th week of gestation (GW). We found that in human embryos, Prox1 is expressed in the cardinal veins at Carnegie stage (CS)12, initiating the emergence of LECs. Subsequently, LECs budding from the cardinal veins begin to express VEGFR3 at CS13. LYVE1 and PDPN are expressed in lymph sacs at CS16. A valve structure is formed on the lymphatic side of the junction between the lymph sac and the cardinal veins at CS18, creating a boundary between the cardinal veins and the lymphatic vessels. On the other hand, when observing in each organ (heart, lower jaw, lungs, mesentery, kidney, and thoracic duct), the development of lymphatic vessels varies.

Our study is the first to clarify the lymphatic vessel development in humans. LECs are preliminary derived from embryonic veins. On the other hand, the process of lymphatic development in each organ is spatiotemporally diverse, which may be due to differences in their developmental processes or cellular origins. These findings indicate that the development of lymphatic vessels is similar between humans and other species. Our research offers essential insights into the evolution and phylogeny of lymphatic vessels, and may also illuminate the pathogenesis of lymphatic-related diseases, which include lymphedema, obesity, cardiovascular disorders, Crohn’s disease, and congenital lymphatic disease, such as lymphatic malformation.

## Results

### Evaluation of marker expression in human fetal lymphatic endothelial cells

To explore the development of human lymphatic vessels, we tested antibodies against Prox1, VEGFR3, and Lymphatic Vessel Endothelial Hyaluronan Receptor-1 (LYVE1), all of which are commonly employed in the study of murine lymphatic vessel development. Additionally, we incorporated D2-40, a monoclonal antibody that specifically interacts with the glycoprotein Podoplanin (PDPN). This antibody is frequently used in human pathological diagnoses for staining to adult lymphatic vessels. We also employed Platelet Endothelial Cell Adhesion Molecule-1 (PECAM) antibodies as a marker for endothelial cells. By GW9, we confirmed that most organs had developed sufficiently to resemble those found in adults. We proceeded with the fluorescent immunostaining and enzyme-antibody method, followed by 3,3′-Diaminobenzidine (DAB) color development in this GW9 fetus and verified the expression of all marker proteins in LECs within jugular lymph sacs (**Supplemental Figure 1A-H**). Additionally, the specificity of the staining was confirmed with controls using only the secondary antibodies (**Supplemental Figure 1I-L**). From these observations, it became evident that these molecules are also expressed in lymphatic vessels in human fetuses.

### The emergence of lymphatic endothelial cells begins in the embryonic cardinal veins in human embryos

Next, we analyzed the development of lymphatic vessels using embryos at various stages. In CS 11, when the formation of the precardinal vein occurred, we could not identify Prox1 expression in the precardinal vein (**Supplemental Figure 2A-C’**). At CS12, we performed immunostaining using PECAM, Prox1, and Chicken ovalbumin upstream promoter– transcription factor II (COUP-TFII) antibodies. COUP-TFII, a member of the nuclear receptor superfamily, is necessary for the activation of Prox1 in the cardinal veins in mice (28, 29). We identified Prox1^+^/PECAM^+^/COUP-TF2^+^ and Prox1^+^/PECAM^+^/COUP-TF2^-^ LECs in and around the anterior cardinal veins (ACVs) (**Figure 1A-I**). At this stage, LYVE1 expression was not observed in the ACVs or its surroundings, whereas VEGFR3 was expressed in the ACVs (**Figure 1C, D, G, H**). By CS13, Prox1^+^/PECAM^+^/Coup-TF2^+^ or Prox1^+^/PECAM^+^/COUP-TF2^-^ LECs formed capillary lymphatics extending towards the posterior of the body (**Figure 1J-K’’’**). While these capillary lymphatics expressed VEGFR3 (**Figure 1L-M’’’**), they did not express LYVE1 (**Figure 1N-O’’’**).

**Figure 1.**
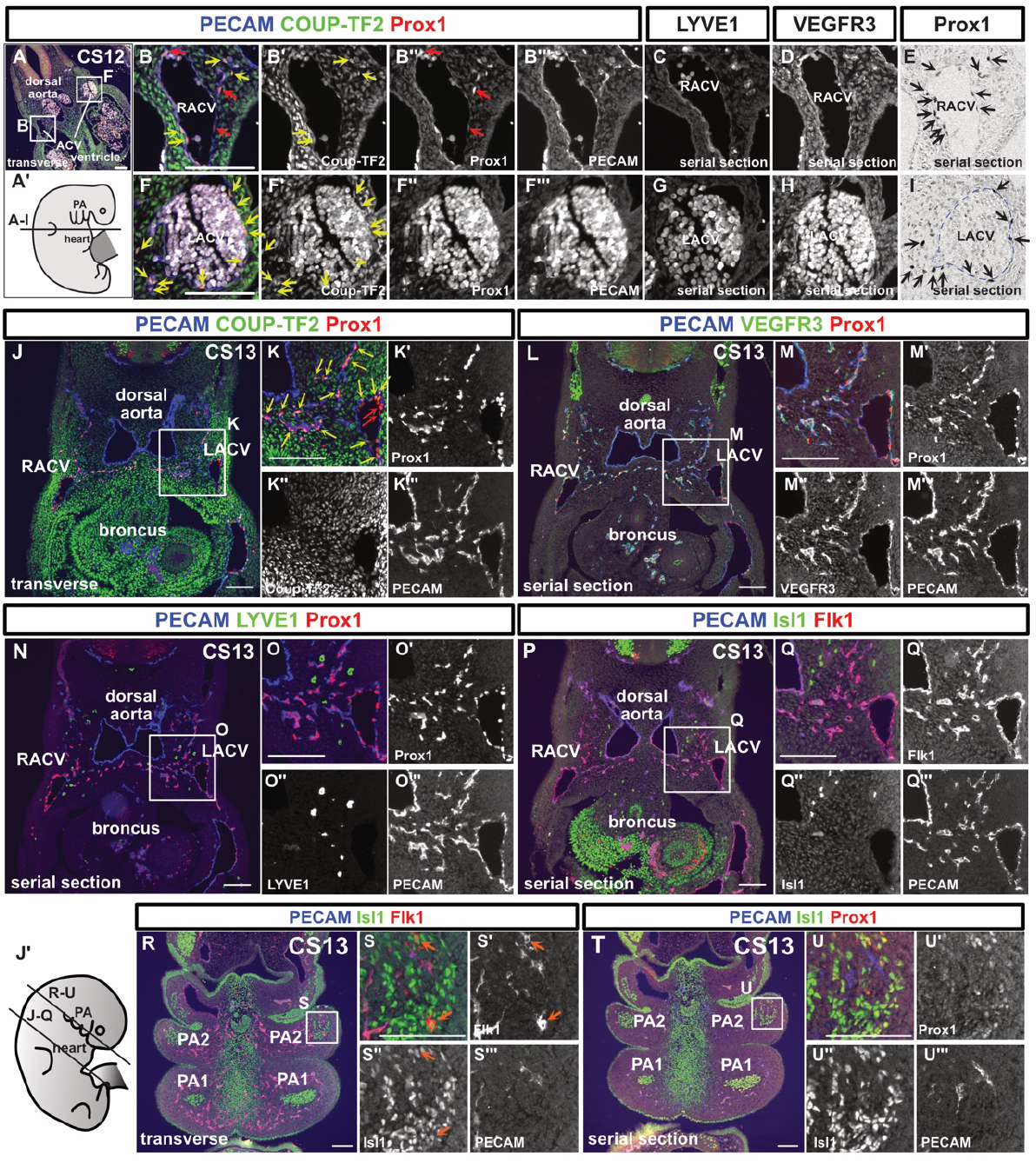
Lymphatic endothelial cells originate from the cardinal veins in human embryos. (A-I) Immunostaining of transverse sections using the indicated antibodies in a CS12 embryo and schematic representation of the locations of the sections; We detected Prox1^+^/PECAM^+^/Coup-TF2^-^ cells (red arrows) and Prox1^+^/PECAM^+^/Coup-TF2^+^ cells (yellow arrows: 78.9% of PECAM^+^/Prox1^+^ cells were Coup-TF2^+^, n=1) in and around the ACVs. On average, there were 18.25 nuclei per cross-section of the CV; of these, 4.5 were Prox1^-^ /PECAM^+^ blood endothelial cells (BECs), and 13.75 were Prox1^+^/PECAM^+^ LECs. Therefore, BECs constituted 24.7%, and LECs constituted 75.3%. There were an average of 9.75 Prox1^+^/PECAM^+^ cells located externally to the CV. (E, I) Enzyme antibody method; Prox1^+^ cells were present around the ACVs (black arrows). These Prox1^+^/PECAM^+^ cells were localized around the cardinal veins in the CS12 embryos. (J-U’’’) Fluorescent immunostaining of transverse sections using the indicated antibodies in a CS13 embryo and a schematic representation of the positions of the sections; The PECAM^+^/Prox1^+^ LECs budding from the ACVs consisted of Coup-TF2-expressing LECs (yellow arrows: 82.8±2.18% of the PECAM^+^/Prox1^+^ cells were Coup-TF2^+^, n=2) and Coup-TF2^-^ LECs. PECAM^-^/Isl1^+^/Flk1^+^cardiovascular progenitor cells were observed in the pharyngeal arch mesoderm (orange arrow). CV, cardinal vein; RACV, right anterior cardinal vein; LACV, left anterior cardinal vein; PA1, first pharyngeal arch; PA2, second pharyngeal arch; scale bar, 100 μm (A, B, F, J, K, L, M, N, O, P, Q, R, S, T, and U)

We previously reported that the lymphatic vessels in the head and neck region originate from *Islet1* (*Isl1*)^+^ LECs located in the pharyngeal arch mesodermal regions(17, 30). Additionally, it was suggested that these LECs may be derived from multipotent Isl1^+^/Flk1^+^ cardiovascular progenitors(31). Based on these findings, we performed immunostaining of Isl1, Flk1, PECAM, and Prox1 to identify LECs and their progenitor cells originating from the pharyngeal arches. We did not detect any Isl1^+^ endothelial cells in and around the cardinal veins (**Figure 1P-Q’’’**). Although, we detected a small number of Flk1^+^/Isl1^+^/PECAM^-^ cells within the second pharyngeal arch mesoderm, we could not find any cells that expressed both Prox1 and Isl1 (**Figure 1R-U’’’**).

At CS14, we observed more Flk1^+^/Is1^+^/PECAM^-^ cells in the second pharyngeal arch (**Supplemental Figure 3A-C’’’**). However, we did not find any cells co-expressed Isl1 and Prox1 (**Supplemental Figure 3D-E’’’**). Around the ACVs, LECs partially formed luminal structures at CS14 (**Supplemental Figure 3F-J**). In the CS15 embryo, we identified a few Prox1^+^ cells and PECAM^+^ cells within the second pharyngeal arch (**Supplemental Figure 3K, K’, P, and P’**). However, due to poor fixation and excessive autofluorescence in multi-fluorescence staining, we were unable to determine whether PECAM and Prox1 were co-expressed in the pharyngeal arches. VEGFR3^+^ cells were present in the second pharyngeal arch, but for the same reason, we could not determine whether they were expressed in LECs (**Supplemental Figure 3M and M’**). The Expression of LYVE1 and PDPN was not observed in the pharyngeal arches (**Supplemental Figure 3N-O’**). At CS16, cells co-expressing Prox1 and Isl1 were not observed in the lower jaw or the cardiac outflow tract regions (**Supplemental Figure 3Q-S’’’**). Additionally, at GW9, Flk1 expression was detected in the cervical lymph sac, but Isl1 expression was not (**Supplemental Figure 1M and N**).

In summary, LECs arise from the ACVs at CS12. We identified Flk1^+^/Isl1^+^/PECAM^-^ cells in the second pharyngeal arch, but we were unable to identify any cells co-expressing Isl1 and Prox1 at CS13, CS14, and CS16. Therefore, we have concluded that in human embryos LECs first originate from the ACVs.

### Lympho-venous valves are formed between lymph sacs and the cardinal veins

At CS16, the large initial lymphatic vessels (lymph sacs) around the ACVs were formed and they expressed LYVE1 and PDPN as well as VEGFR3 and Prox1 (**Figure 2A-D**). By CS18, the connection between the ACVs and lymph sacs was observed and on lymph sacs side, valve-like protrusions began to appear (**Figure 2E-M**). In the junction where the cardinal veins and lymphatic sacs merged, no VEGFR3, Prox1, LYVE1, or PDPN expression was seen on the venous side, clearly delineating the boundary between the lymphatic vessels and veins (**Figure 2F-L**). Notably, while the venous side maintained a smooth lumen, the lymphatic side appeared distorted, facilitating morphological distinction (**Figure 2E-M**).

**Figure 2.**
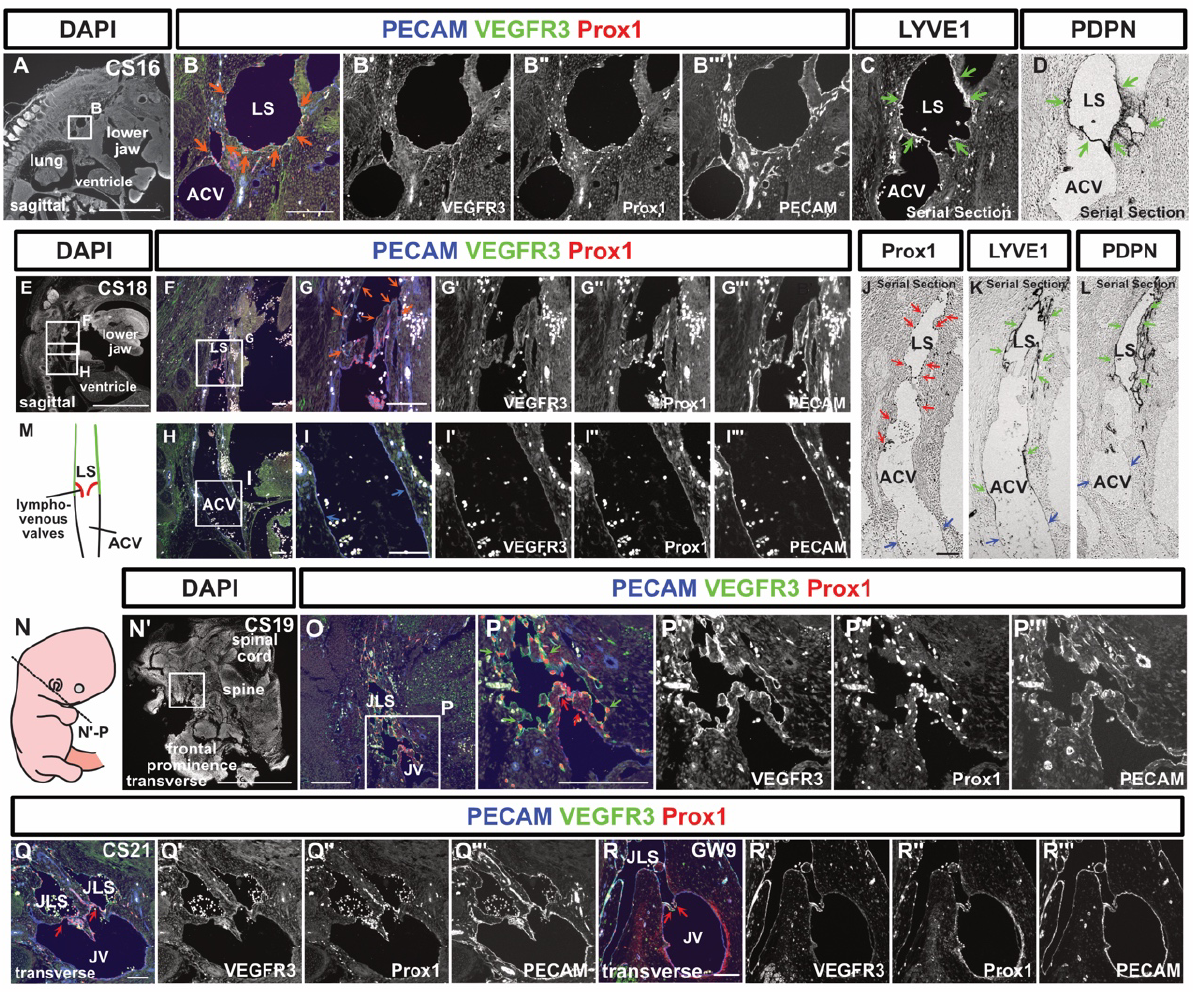
Lymph sacs express LYVE1 and PDPN and establish connections with the anterior cardinal veins to form lympho-venous valves. (A-D) Immunostaining of sagittal sections from a CS16 embryo using the indicated antibodies; (B-C) Fluorescent immunostaining; (D) Immunostaining using the enzyme-antibody method. At CS16, lymph sacs emerged, which exhibited Prox1, VEGFR3, and PECAM expression (orange arrows). (C, D) The expression of LYVE1 and PDPN began within these lymph sacs at this stage (green arrows). (E-L) Immunostaining of sagittal sections from a CS18 embryo using the indicated antibodies; (E-I’’’) Fluorescent immunostaining; (J-L) The enzyme-antibody method; (E-I’’’) Lymph sacs expressing PECAM, VEGFR3, and Prox1 (orange arrows) formed connections with the ACVs, which expressed PECAM (blue arrows). (J-L) While Prox1 (red arrows), LYVE1, and PDPN (green arrows) were expressed in the lymph sacs, these markers were absent from the ACVs (blue arrows). There were 4 CS18 stage embryos, but only one included lymph sacs. (M) Schematic representation of the relationship between the ACVs and lymph sacs in a CS18 embryo. The lymph sacs and the ACVs were continuous, featuring a valve structure on the lymph sac side. (N-P’’’) Transverse sections and schematic representation of the positions of the sections in a CS19 embryo; Fluorescent immunostaining of PECAM, Prox1, and VEGFR3 was conducted. VEGFR3 was expressed in lymph sacs at CS19 (green arrows). The LVVs were overlaid with Prox1^+^ cells (red arrows) (LVVs were identifiable in 1 of 5 embryos). (Q-R’’’) Transverse sections of a CS21 embryo and a GW9 fetus, highlighting the areas containing LVVs; As was the case at CS19, LVVs were visible (red arrows). (At CS21, LVVs were identified in 1 out of 2 embryos; at GW9, LVVs were identified in 1 out of 4 fetuses). LS, lymph sacs; ACV, anterior cardinal vein; JLS, jugular lymph sacs; JV, jugular vein; scale bars, 1 mm (A, E, and N’) or 100 μm (B, F, G, H, I, J, O, P, Q, and R)

Similar findings were observed at stages CS19 and 21. At the junction between jugular lymph sacs and the jugular vein, lympho-venous valves (LVVs) were formed, and VEGFR3 expression was visible on the lymph sac side, but absent from the venous side (**Figure 2N-Q’’’**). On both the lymph sac and venous sides, the valve leaflets were covered by Prox1^+^ LVV endothelial cells (**Figure 2N-Q’’’**). The LVVs, which were initially irregularly shaped at CS18, gradually assumed a regular form as development progressed; i.e., by GW9 a bicuspid valve had developed (**Figure 2R-R’’’**).

### The development of lymphatic vessels varies among organs

Consequently, we conducted an analysis of lymphatic vessel development across various organs.

#### The cardiac lymphatic vessels

At CS16, capillary lymphatic vessels and isolated LECs in the wall of the aorta were observed (**Figure 3A**). This lymphatic network was composed of PECAM^+^/Prox1^+^/VEGFR3^+^ or PECAM^+^/Prox1^+^/VEGFR3^-^ LECs, which did not express LYVE1 or PDPN (**Figure 3A-E**). As development progressed, tubular lymphatic vessels were identifiable at CS23 (**Figure 3F-O**). Throughout this process, the initially mesh-like capillary lymphatics undergo progressive remodeling to establish lumen-bearing vessels. Consequently, while the density of LECs per unit area remains relatively stable, there is an increase in the number of lymphatic vessels possessing distinct luminal structures (**Figure 3N and O**). These lymphatic vessels expressed LYVE1 and PDPN at CS23 and GW9 (**Figure 3L and M, Supplemental Figure 4X’ and Y’**). By GW9, lymphatic vessels were identifiable on the epicardial side (**Figure 3P and Q**). Lymphatic vessels were also distributed around the coronary arteries (**Figure 3P-R**). No lymphatic vessels were seen in the myocardium or endocardium during the examined periods (**Figure 3S**).

**Figure 3.**
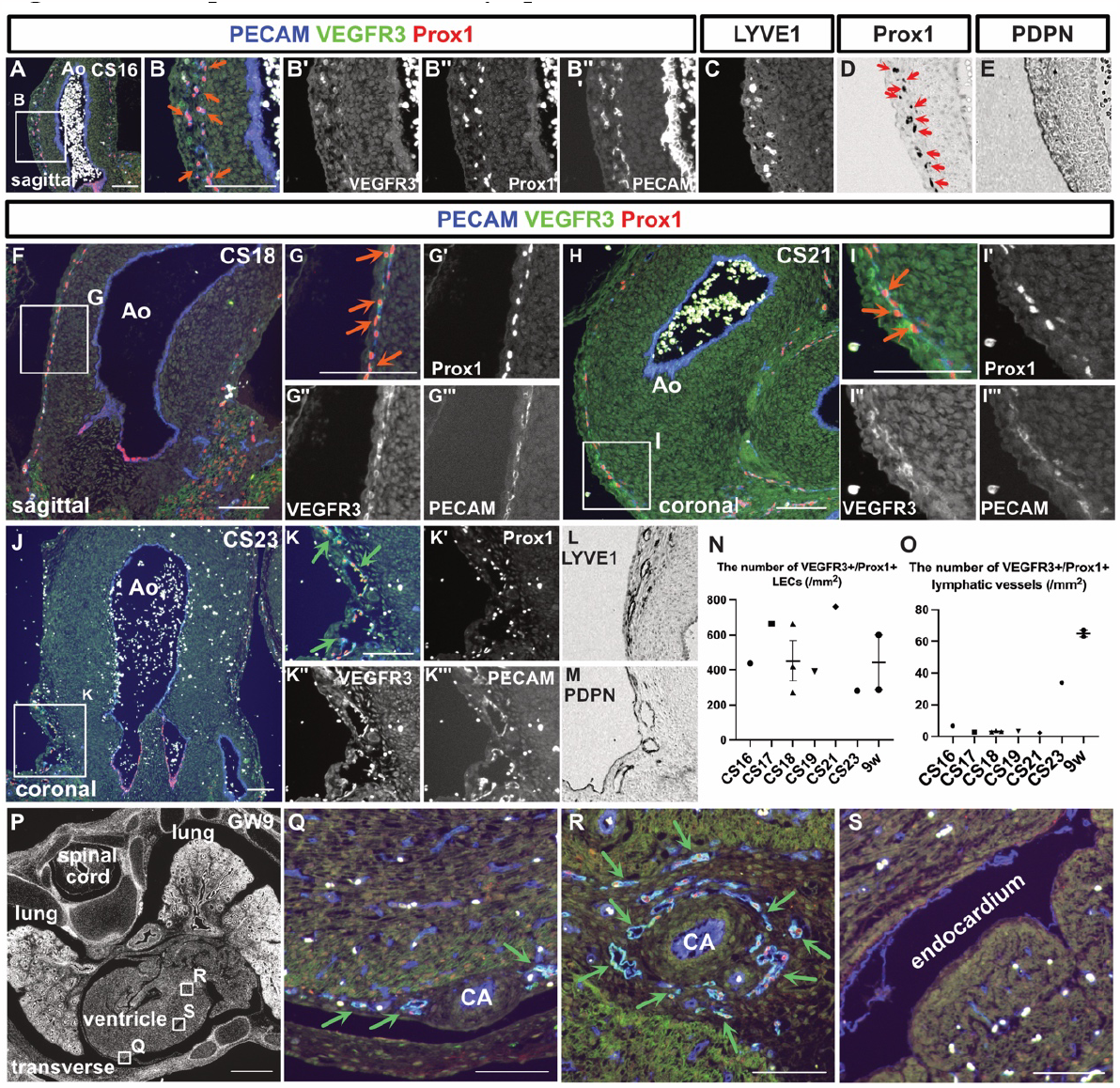
Development of the cardiac lymphatic vessels. (A-E) Immunostaining of sagittal sections from a CS16 embryo using the indicated antibodies; (A-C) Fluorescent immunostaining; (D, E) Enzyme-antibody method, followed by DAB color development; PECAM^+^/Prox1^+^/VEGFR3^+^ LECs were present in the aortic wall (orange arrows). (D) Prox1^+^ cells were also identified by the enzyme-antibody method (red arrows). (F-I’’’) Immunostaining of sections from CS18 and CS21 embryos using the indicated antibodies. Consistent with the CS16 embryo, PECAM^+^/Prox1^+^/VEGFR3^+^ LECs could be identified in the aortic wall (orange arrows). (J-M) Immunostaining of coronal sections from a CS23 embryo using the indicated antibodies; (J-K’’’) Fluorescent immunostaining; (L, M) Enzyme-antibody method, followed by DAB color development; LYVE1^+^ and PDPN^+^ lymphatic vessels were identified around the aorta (green arrows). (N, O) Changes in the number of PECAM^+^/Prox1^+^/VEGFR3^+^ LECs in the aortic wall as development advanced (N); Changes in the number of PECAM^+^/Prox1^+^/VEGFR3^+^ luminal lymphatic vessels as development advanced (O). Each dot represents a single individual. (P-S) Fluorescent immunostaining of PECAM, Prox1, and VEGFR3 in transverse sections of a GW9 fetus. By GW9, lymphatic vessels could be seen around the coronary arteries in the epicardium (green arrows). Ao, aorta; CA, coronary artery; scale bars, 1 mm (P) or 100 μm (A, B, F, G, H, I, J, K, Q, R, and S).

#### Lung lymphatic vessels

At stage CS16, only a small number of PECAM^+^/Prox1^+^/VEGFR3^+^ LECs could be detected in the lung parenchyma (**Figure 4A**). However, as development progressed increases in the number of LECs and luminal lymphatic vessels were evident (**Figure 4A-J**). At CS 23, lymphatic vessels were distributed around the trachea, which became apparent by GW9 (**Figure 4G and H**). PDPN and LYVE1 expression were detected in the lymphatic vessels in the lung at GW9 (**Supplemental Figure 4X and Y**).

**Figure 4.**
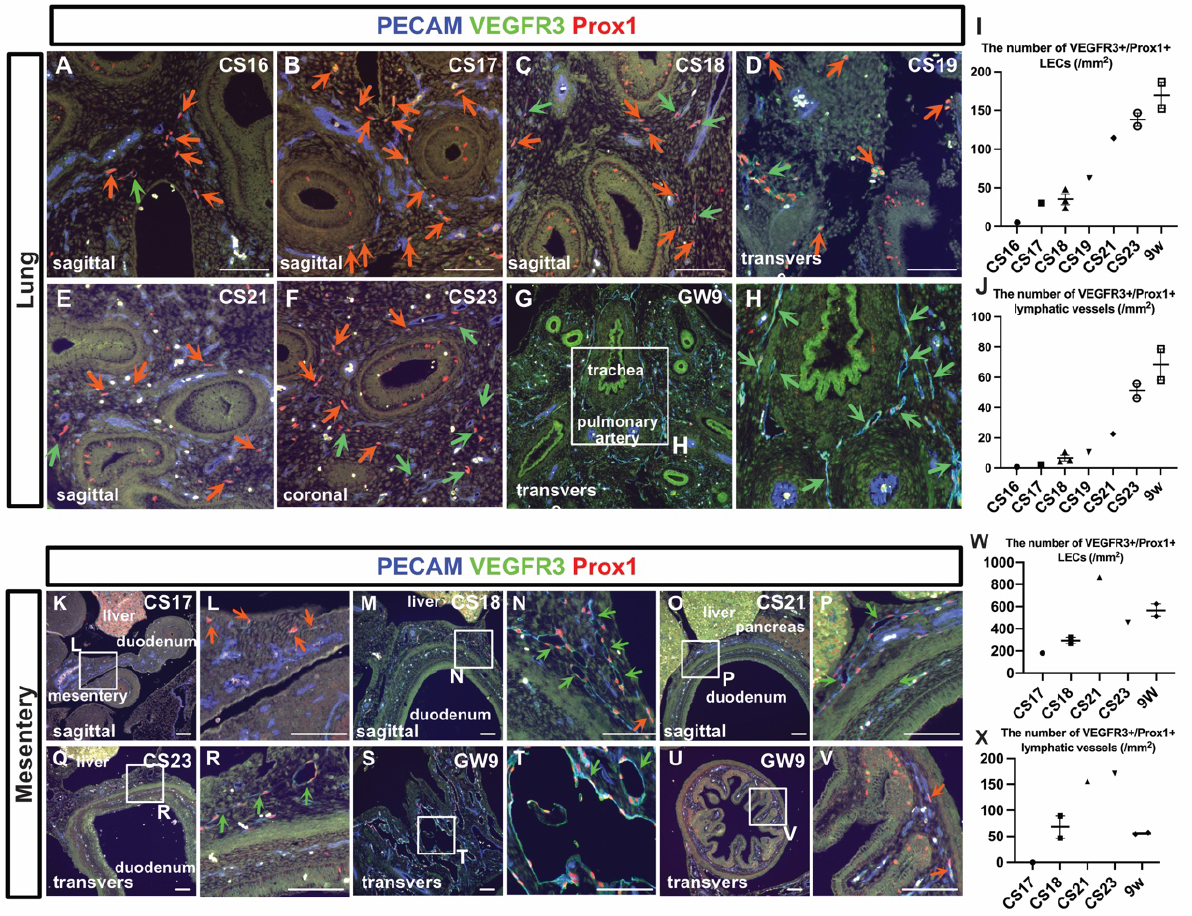
Development of the lung lymphatic vessels and mesenteric lymphatic vessels. (A-H) Immunostaining of PECAM, Prox1, and VEGFR3 in the lungs at each stage; The orange arrows indicate isolated or small clusters of LECs. The green arrows indicate lymphatic vessels that had formed luminal structures. (I, J) Changes in the number of PECAM^+^/Prox1^+^/VEGFR3^+^ LECs in the aortic wall as development advanced (I). Changes in the number of PECAM^+^/Prox1^+^/VEGFR3^+^ luminal lymphatic vessels as development advanced (J). Each dot represents a single individual. (K-V) Immunostaining of PECAM, Prox1, and VEGFR3 in the intestine and mesentery at each indicated stage. The orange arrows indicate isolated or small clusters of LECs that had not formed luminal structures, while the green arrows indicate lymphatic vessels that had formed luminal structures. (W, X) Changes in the number of PECAM^+^/Prox1^+^/VEGFR3^+^ LECs in the aortic wall as development advanced (W). Changes in the number of PECAM^+^/Prox1^+^/VEGFR3^+^ luminal lymphatic vessels as development advanced (X). Each dot represents a single individual. Scale bars, 100 μm (A-H and K-V).

**Figure 5.**
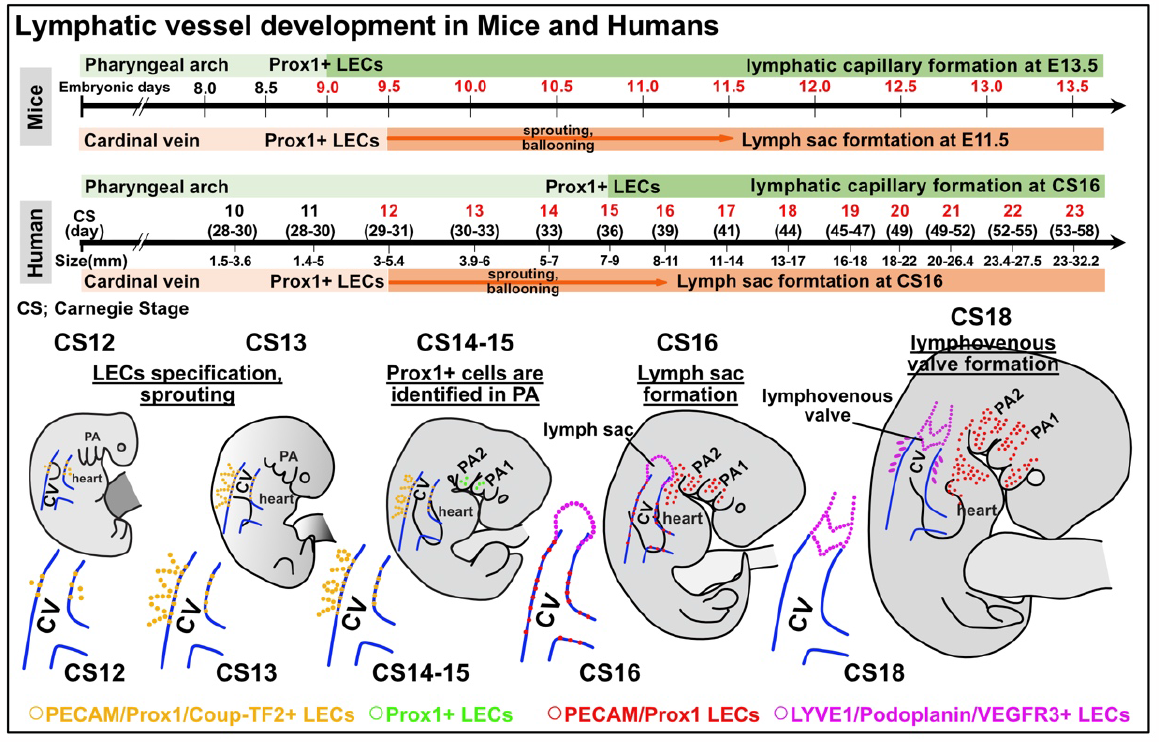
Comparative overview of early lymphatic vessel development in mouse and human embryos. In mouse embryos, Prox1 expression is initiated in the cardiac pharyngeal mesoderm from E9.0 to E9.5, leading to the distribution of LECs in the head and neck, mediastinal, and cardiac outflow regions. By E13.5, these LECs start forming capillary lymphatics, which further develop into larger lymphatic vessels. Prox1 expression in the cardinal vein starts at E9.5, with the lymph sac forming post-sprouting by E11.5. In contrast, for human embryos, Prox1 expression in the pharyngeal arch area is first noted at CS15. Following this, LECs in the mandibular and cardiac outflow tract regions slowly establish a capillary lymphatic network, which by CS23 progresses to clearly defined lymphatic vessels with luminal structures. Prox1 expression in the cardinal vein begins at CS12, with the lymph sac being discernible by CS16. By CS18, the lympho-venous valve is identifiable between the lymph sac and the ACV. Across both embryos, early expression of transcription factors such as Prox1 and Coup-TF2 is observed, with the subsequent expression of VEGFR3, LYVE1, and Podoplanin. CS, Carnegie stage; ACV, the anterior cardinal veins.

#### Mesenteric and intestinal lymphatic vessels

At stage CS17, PECAM^+^/Prox1^+^/VEGFR3^+^ LECs were identified within the mesentery of the midgut, although no luminal lymphatic vessels were detectable (**Figure 4K and L**). By stage CS18, the LECs in the mesentery had started forming luminal structures (**Figure 4M and N**). At stage CS21, although lymphatic vessels were evident in the mesentery, only PECAM^+^ blood vessels could be observed in the lamina propria of the intestine (**Figure 4O and P**). This was still the case at CS23 (**Figure 4Q and R**). By GW9, enlarged lymphatic vessels were seen in the mesentery (**Figure 4S and T**), and a small number of PECAM^+^/Prox1^+^/VEGFR3^+^ LECs were present in the lamina propria of the intestine (**Figure 4U and V**). In summary, starting from CS17, LECs could be identified in the mesentery, and their numbers gradually increased, eventually resulting in the formation of large-diameter lymphatic vessels in the mesentery (**Figures 4W and X**). PDPN and LYVE1 expression were detected in the lymphatic vessels within the mesentery and intestinal wall at GW9 (**Supplemental Figure 4X’’’, X’’’’, Y’’’, and Y’’’’**).

#### Lymphatic vessel development in the lower jaw

Similar to the findings for cardiac lymphatic vessels, in mouse embryos roughly 70-80% of the lymphatic vessels in the lower jaw originated from the cardiopharyngeal mesoderm (17). In line with this, the developmental pattern was resemble that of the lymphatic vessels surrounding the aorta. At stage CS16, PECAM^+^/Prox1^+^/VEGFR3^+^ lymphatic vessels formed a small number of luminal structures without LYVE1 or PDPN expression (**Supplemental Figure 4A-C’’’**). The majority of LECs were present either as isolated or a few linked. This pattern persists at least until CS22 (**Supplemental Figure 4A-Q**). At GW 9, tubular lymphatic vessels are present in the subcutaneous tissue, expressing LYVE1 and PDPN (**Supplemental Figures 4X’’’’’ and Y’’’’’**).

#### Kidney lymphatic vessel development

In mouse embryos, lymphatic vessels develop from the renal hilum within the metanephros, which matures into the adult kidney, and they gradually extend throughout the entirety of the kidney (32). At stage CS23, we observed a few LECs in the renal hilum of metanephros. At GW9, increased lymphatic vessels were detected around blood vessels in the renal hilum eith the expression of LYVE1 and PDPN(**Supplemental Figure 4R-U, 4X’’ and Y’’**).

#### Thoracic duct development

The thoracic duct is the largest lymphatic vessel in the human body, distributed along the posterior peritoneum, aorta, and esophagus. At GW9, large-diameter lymphatic vessels were observed around the aorta, which may eventually combine to form the future thoracic duct (**Supplemental Figure 4V and W**).

## Discussion

In this study, we examined the lymphatic vessel development process using human embryos. We identified antibodies that could be used in human embryos and fetuses. Using these antibodies, we identified endothelial cells expressing Prox1 and Coup-TF2, which are necessary for the production of early LECs, in the cardinal vein at CS12. On the other hand, while we tried to identify *Isl1*^*+*^ LECs derived from the cardiopharyngeal mesoderm that we reported in mice(17), we were unable to find any LECs that simultaneously expressed both Isl1 and Prox1 at CS13, 14, and 16. This may have been due to the disappearance of Isl1 expression before Prox1 expression. However, we observed Flk1^+^/Isl1^+^/PECAM^-^ cells in the pharyngeal arch mesodermal region, and such cardiovascular progenitor cells(31) may form lymphatic vessels in the head and neck region by eventually expressing Prox1.

LECs budding from the cardinal vein gradually aligned and formed lymph sacs around the cardinal vein at CS13 to 16. LECs did not express LYVE1 and PDPN from CS12 to CS15, however, when they formed lymph sacs at CS16, they exhibited LYVE1 and PDPN expression, as well as Prox1 and VEGFR3. This marker expression patterns aligned with our previous works regarding to mice cardiac lymphatic vessel development, showing that LECs first express Prox1, then VEGFR3, followed by the expression of LYVE1(4, 30). We also confirmed that LVVs formed at the junction between the lymph sacs and the cardinal veins at CS18. LVVs initially showed an irregular shape, but gradually formed smooth bicuspid valves around GW9. The proper formation of LVVs is critical as a cause of primary lymphedema in humans(33). Therefore, the understanding of the normal LVVs development from this study may help to elucidate the pathology of primary lymphedema caused by genetic mutations(1).

Focusing on the lymphatic vessel development of each organ, for example, the lymphatic vessels in the lower jaw and cardiac outflow tract, which we reported to be derived from the cardiopharyngeal mesoderm, did not exhibit noticeable luminal formation at CS16-21. At CS22, many LECs still capillary lymphatics. In the heart, LECs gradually gathered together, luminal formation accelerated, and PDPN and LYVE1 expression were observed at CS23. In the lower jaw, the expression of PDPN and LYVE1 and the formation of luminal structures were confirmed at GW9. In the lungs, the number of LECs increased and tubular structures started to form as development progressed. By GW9, we were able to confirm the presence of lymphatics around the bronchi, in a similar distribution to that seen in adults. At CS17, isolated LECs were observed within the mesentery, which was continuous with the posterior abdominal wall. At this stage, although lymph sacs had developed within the posterior abdominal wall (data not shown), no continuity with the isolated LECs was observed. As development progressed, the mesenteric LECs formed large lymphatic vessels. However, the presence of LECs in the intestinal wall was not observed until GW9. In the kidney, considering that only a few LECs were observed around CS23, the lymphatics may begin to develop around CS23 in human embryos. Also, in order to consider the development of the thoracic duct, we confirmed the presence of large-diameter lymphatic vessels around the aorta and esophagus at GW9. However, the course and shape of the thoracic duct vary greatly among individuals, so a more detailed analysis will likely be needed to determine how it develops. Given that genetic cell lineage analysis in humans requires significant challenges, combining developmental analysis with multi-omic analysis, such as single-cell RNA-seq analysis, could potentially clarify the details of organ-specific lymphatic vessel development.

Although recent research has shown that the meningeal lymphatic vessels play a crucial role in the pathophysiology of the central nervous system (CNS) (1), we were unable to confirm the presence of the lymphatic vessels in the brain and spinal cord. Indeed, in mice the CNS lymphatic vessels develop after birth(34).

The regulation of Prox1 expression requires transcription factors, such as Coup-TF2(28) and Sox18(35) to the promoter region. We confirmed the expression of Coup-TF2 in the cardinal vein and surrounding LECs, however the expression of Sox18 in the cardinal vein and LECs was not observed in human embryos at CS13 (data not shown). This might be due to the expression timing of Sox18 is more restricted in the cardinal vein in humans. In the process of LVVs formation, in addition to Prox1, the upregulation of FOXC2 and GATA2 leads to the acquisition of the differentiation fate towards LVV endothelial cells(36). We attempted to elucidate the expression patterns of these molecules, but the number of sections containing LVVs in a single individual was limited; therefore, we were not able to analyze these expression patterns.

In summary, this study clarified the process underlying early lymphatic vessel formation in human embryos. LECs originate from embryonic veins and form lymph sacs, a process which has been clarified in mice and zebrafish and is now confirmed to also occur in humans. On the other hand, the lymphatic vessel development processes for each organ vary in terms of speed and marker expression, possibly due to differences in cellular origin and signaling. This study is important for elucidating the evolutionarily conserved processes of lymphatic vessel formation and for aiding the extrapolation of findings from animals to humans.

## Materials and Methods

### Tissue collection and ethical considerations

For this study, 31 preserved human embryos in organogenesis period and 3 fetuses ranging from CS8 to GW9 were analyzed. The samples used were collected for pathological examination between January 2000 and October 2021 and had been preserved for the purpose of pathological diagnosis. The use of human samples was approved by the ethics committee of Mie University Hospital (approval number: H2021-228). No informed consent (written or verbal) was obtained, since this was not deemed necessary by the ethics committee. The staging of the embryos and fetuses used in the experiments was done using a combination of the Carnegie stage (CS) (37) and clinical information: 1) menstrual weeks, 2) morphology, 3) length (crown-rump), and 4) anatomical features (we also referred to the detailed anatomical information on this website: https://www.ehd.org/virtual-human-embryo/). Detailed information regarding each sample is presented in Table 1. The sex of each sample was not determined, with the exception of one case of miscarriage. In this particular case, chromosomal analysis verified the absence of any karyotypic abnormalities. There were no malformations observed in any of the embryos or fetuses. Nevertheless, for the remaining embryos, there is a possibility that developmental defects or mutations could lead to abnormalities in the lymphatic vessels.

**Table 1.** Information of human embryos and fetuses.

|  | Number and characteristics of embryos and fetuses. | Cause and additional information of embryos and fetuses |
| --- | --- | --- |
| CS9-10 | N=1. The heart tube formation. | Miscarriage. 5 weeks from LMP. |
| CS10 | N=2. The neural tube formation. | Miscarriage (N=2). Details unknown. The patients bring miscarriage specimens. |
| CS11 | N=1. GL is 1.4 to 5 mm. Precardinal veins formation. | Tubal pregnancy. GL was 5mm (transvaginal ultrasound). 5 weeks from LMP. |
| CS12 | N=1. GL is 3 to 5.4 mm. Trabecular formation in the heart. | The samples was taken by emergency surgery due to tubal pregnancy. |
| CS13 | N=2. GL is 3.9 to 6 mm. The lung buds formation. | The samples was taken by emergency surgery due to tubal pregnancy. |
| CS14 | N=2. GL is 5 to 7mm. | Miscarriage (N=2). 6 weeks from LMP. GL was 6 mm (N=1, transvaginal ultrasound). The size of the other was unknown. |
| CS15 | N=1. GL is 7 to 9 mm. The lens pit formation. | Miscarriage. 7 weeks from LMP. GL was 5.5 mm (transvaginal ultrasound). |
| CS16 | N=1. GL is 8 to 11 mm. The retinal pigment is distinct. | The samples was taken by emergency surgery due to tubal pregnancy. |
| CS17 | N=1. GL is 11 to 14 mm. | Tubal pregnancy. 8weeks from LMP. GL was 14.0 mm (transvaginal ultrasound). |
| CS18 | N=4. GL is 13 to 17 mm. Posterior cardinal vein has disappeared. | Artificial abortion (N=3), tubal pregnancy (N=1). 8 weeks from LMP. GL was 14.0, 15.0, and 16.5 mm (transvaginal ultrasound). The other was unknown. |
| CS19 | N=5. Embryos have a greatest length of 16 to 18 mm. | Artificial abortion (N=5), tubal pregnancy (N=1). 8 weeks from LMP. GL was 17 mm (N=1, the actual size of the embryo). The others was un known. |
| CS20 | N=1. The CRL is around 17.0-19.3 mm. | Artificial abortion. 8 weeks from LMP. CRL was 18.7 mm (transvaginal ultrasound) |
| CS21 | N=2. The CRL is around 18.8-23.0 mm. | Miscarriage and tubal pregnancy. 9 weeks from LMP. CRL=23 mm (N=1, transvaginal ultrasound). The other was unknown. |
| CS22 | N=3. The CRL is around 22.0-24.0 mm. A few large glomeruli are present in the kidney. | Artificial abortion (N=3). 10 weeks from LMP. No ultrasound information was available. |
| CS23 | N=4. The CRL is around 23.5-28.0mm. Most organs are fully formed. | Artificial abortion (N=4). 10-11 weeks from LMP. CRL=29.4mm (N=1). The others was unknown. |
| GW9 | N=3. The CRL is around 25-45 mm<sup>1</sup>. | Artificial abortion (N=1), miscarriage (N=2). CRL was 40.0 and 39.4mm (transvaginal ultrasound). The actual size was 42mm (N=1). |

GL, Greatest length of the embryo; CRL, The crown rump length. LMP; The last

### Section preparation and screening for lymphatic endothelial cells

The embryos and fetuses were fixed with 10% formalin neutral buffer solution at 4 °C for 1 to 2 days. The specimens embedded in paraffin were cut into 1-µm thick sections until the sample was exhausted. Hematoxylin and eosin (HE)-stained sections were created at 10-µm intervals to confirm the anatomical structures of the specimens. Considering the impact of autofluorescence, all staining procedures were coupled with enzyme immunohistochemistry (IHC) to confirm the specificity of the signals. To screen for lymphatic vessels, immunostaining of Prox1, CD31, and LYVE1 was performed on one out of every ten sections using enzyme IHC. After stage CS16, immunostaining of PDPN was also performed when screening for lymphatic vessels.

### Immunohistochemistry (IHC)

HE staining and IHC were performed using 1-μm thick sections. In IHC, sections were deparaffinized and rehydrated through a series of xylene and ethanol. For the enzyme-antibody method, endogenous peroxidase activity was blocked using 0.3% hydrogen peroxide (H_2_O_2_) in methanol for 20 min. In fluorescent antibody staining, to suppress autofluorescence, samples were incubated in 0.1% sodium borohydride in 0.1M phosphate-buffered saline (PBS) (137 mM NaCl, 2.7 mM KCl, 10 mM Na2HPO4, and 1.8 mM KH2PO4, pH=7.2) for 30 minutes, then rinsed with water, and subsequently incubated for 5 minutes in 0.2M glycine in 0.1M PBS. Antigen retrieval was carried out using a pressure chamber with Tris-EDTA buffer (7.4 mM Tris, 1 mM EDTA-2Na, pH 9.0). Slides were incubated with primary antibodies against Prox1 (11-002, AngioBio, 1:150, RRID:AB_10013720), Prox1 (AF2727, R&D Systems, 1:150, RRID:AB_2170716), LYVE1 (ab14917, abcam, 1:150, RRID:AB_301509), VEGFR-3 (AF349, R&D Systems, 1:100, RRID:AB_355314), Coup-TF2 (EPR18443, abcam, 1:150, RRID:AB_2895604), D2-40 (anti-PDPN) (413151, nichirei biosciences, at its original concentration (ready to use product)), Flk1 (AF357, R&D Systems, 1:150, RRID:AB_355320), Isl1 (PA5-27789, Invitrogen, 1:200, RRID:AB_2545265), and PECAM (M0823, DAKO, 1:100, RRID:AB_2114471). In the enzyme-antibody method, the secondary antibody from the Histofine Simple Stain System (Nichirei biosciences) was incubated with the slides for 1 hour. Peroxidase activity was visualized using DAB-H_2_O_2_. For fluorescent immunostaining, Alexa Fluor-conjugated secondary antibodies (Abcam, 1:400) were subsequently applied. After this, True Black (Biotium) was used according to the manufacturer’s instructions to suppress autofluorescence. Imaging was carried out using a Keyence BZ-X700 microscope. All images were processed using ImageJ software.

### Statistical analysis

The numbers of LECs and lymphatic vessels are represented as the average of two or more immunostained slides. Image processing was conducted using ImageJ (NIH). Data are presented as the mean ± standard error of the mean (SEM).

## Acknowledgments

We thank all of the laboratory members for their helpful discussion and encouragement. We also thank Dr. Hiroki Kurihara (the university of Tokyo) for discussing the contents of the paper. This study was supported in part by Grants-in-Aid for Scientific Research from the Ministry of Education, Culture, Sports, Science, and Technology, Japan (20K17072 and 23K15949 to K.M.); the Japan Foundation for Applied Enzymology (VBIC to K.M.); the Miyata Foundation Bounty for Pediatric Cardiovascular Research (K.M.); the SENSHIN Medical Research Foundation (K.M.); Mochida Memorial Foundation for Medical and Pharmaceutical Research (K.M.); Japan Agency for Medical Research and Development (AMED) under Grant Number 22jm0610079h0001 (K.M.); Takeda Science Foundation (K.M.); Mie University: Research promotion and graduate school reform-related research grant project <Interdisciplinary Collaborative Research Support Project> (K.M.); Mie medical research foundation (K.M.); TERUMO life science foundation (K.M.); Kurozumi medical foundation (K.M.).

## Data sharing and resource availability

Further information and requests for resources and reagents should be directed to and will be fulfilled by the lead contact, Kazuaki Maruyama. All data and materials associated with this study are available in the main text.

