## Supplemental Figure and legends for "Lymphatic vessel development in human embryos"

**Supplemental Figure 1. Multiple lymphatic markers are expressed in fetal lymph sacs.**

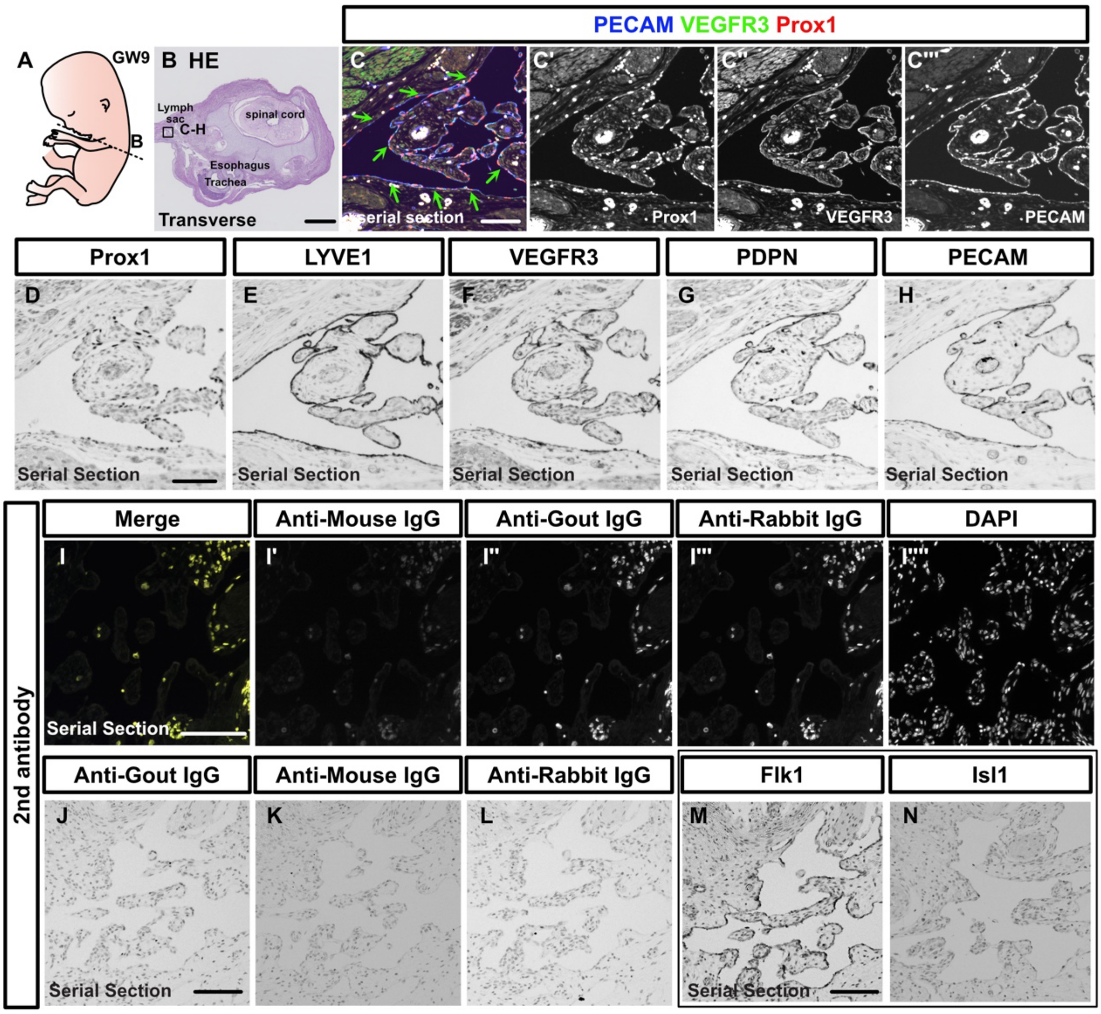


(A-H) Schema showing the positions of the sections in a GW9 fetus (A), and immunostaining of transverse sections (B-H); (C-C''') Fluorescent immunostaining of PECAM, Prox1, and VEGFR3; These markers were expressed in lymph sacs (green arrows). (D-H) Immunostaining of Prox1, LYVE1, VEGFR3, PDPN, and PECAM using the enzyme-antibody method, with color development by DAB. (I-L) Imaging of the secondary antibody-only staining. (M, N) Immunostaining of Flk1, and Isl1 using the enzyme-antibody method, with color development by DAB ; Scale bars, 1 mm (B) or 100 µm (C, D, I, J, and M).

**Supplemental Figure 2. Prox1 expression is not observed in the precardial vein of the CS11 embryo.**

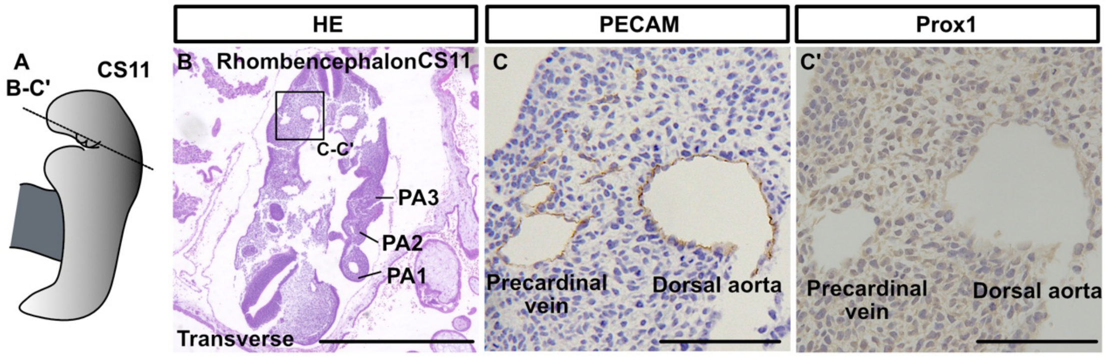


(A-C’) Immunostaining of transverse sections of a CS11 embryo with the indicated antibodies and schema showing a CS11 embryo (n=1). Scale bars, 1 mm (B) or 100 µm (C and C’).

**Supplemental Figure 3. LECs bud from the cardinal veins and form luminal structures.**

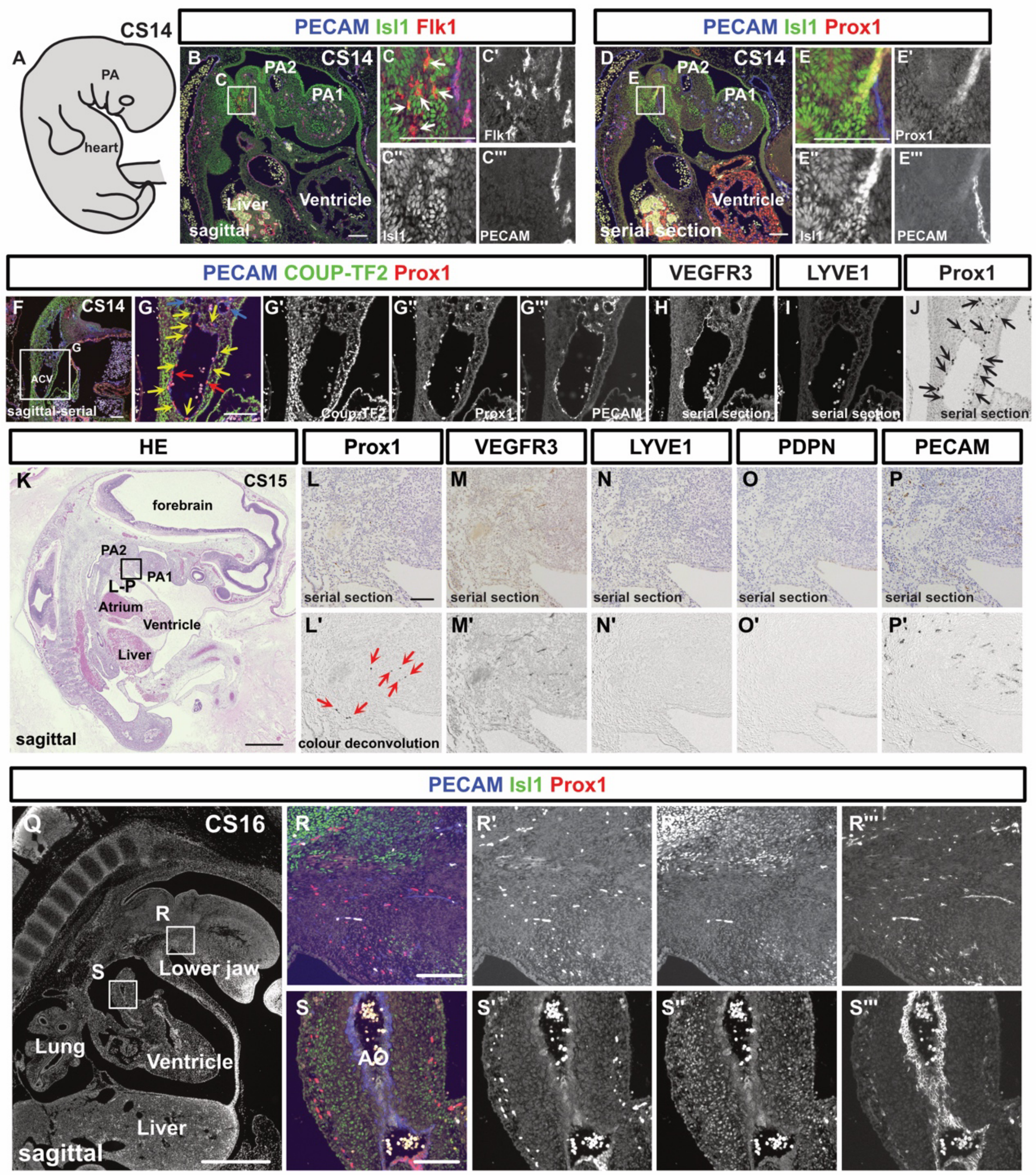


(A-J) Immunostaining of sagittal sections of a CS14 embryo with the indicated antibodies and schema showing the CS14 embryo. (B-C''') Cardiovascular progenitor cells, which were composed of FLK1$^+$/Isl1$^+$/PECAM$^-$ cells, were observed in the second pharyngeal arch (white arrows). (F-G''') PECAM$^+$/Prox1$^+$/Coup-TF2$^+$ cells (yellow arrows: Frequency of Coup-TF2$^+$ cells among PECAM$^+$/Prox1$^+$ cells=44.1%, n=1 [the ACVs could not be identified in another embryo]), and PECAM$^+$/Prox1$^+$/Coup-TF2$^-$ cells (red arrows) were observed in and around the ACVs. (K-P') HE staining of a sagittal section of a CS15 embryo (K) and immunostaining of sagittal sections of the same CS15 embryo with the indicated antibodies (L-P'); At this stage, scattered Prox1$^+$ cells were observed in the pharyngeal arch (red arrows). (Q-S''') Immunostaining of sagittal sections of a CS16 embryo with the indicated antibodies. PA1, first pharyngeal arch; PA2, second pharyngeal arch; ACV, anterior cardinal vein; scale bars, 1 mm (K) or 100 µm (B, C, D, E, F, G, and L).

**Supplemental Figure 4. Development of lymphatic vessels in the lower jaw, kidneys, and thoracic duct.**

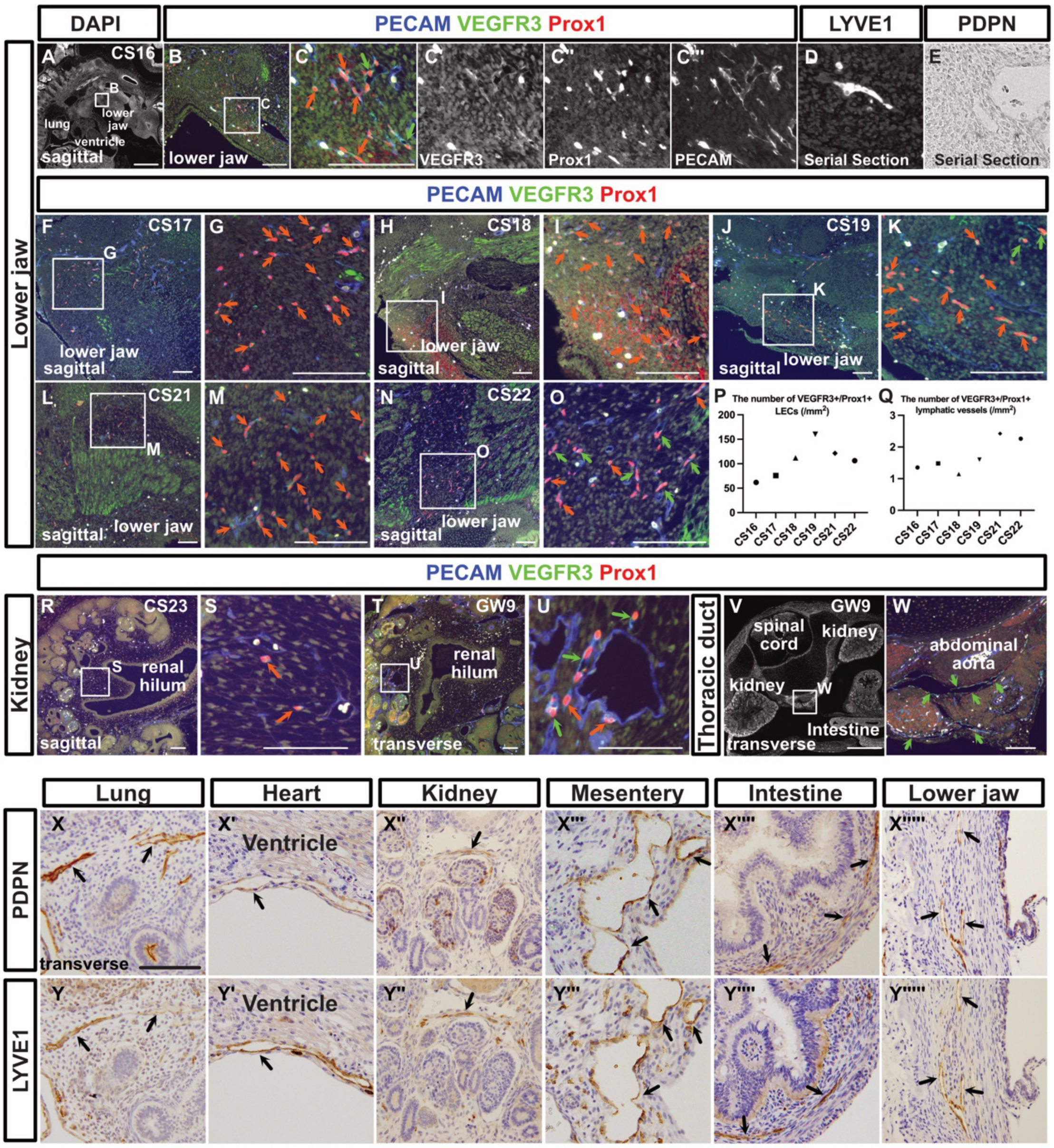


(A-O) Immunostaining of sagittal sections of embryos collected at the described stages with the indicated antibodies; The orange arrows indicate isolated or small clusters of LECs. The green arrows indicate lymphatic vessels that had formed luminal structures. (P, Q) Changes in the number of PECAM$^+$/Prox1$^+$/VEGFR3$^+$ LECs in the aortic wall (P). Changes in the number of PECAM$^+$/Prox1$^+$/VEGFR3$^+$ luminal lymphatic vessels (Q). Each dot represents a single individual. (R-U) Fluorescent immunostaining of PECAM, Prox1, and VEGFR3 in sections from embryos collected at the described stages. The orange arrows indicate isolated or small clusters of LECs. The green arrows indicate lymphatic vessels that had formed luminal structures. (V, W) The green arrows indicate lymphatic vessels surrounding the aorta at GW9. (X, Y) Immunostaining for PDPN and LYVE1 in the lung, heart, kidney, mesentery, intestine, and lower jaw. Scale bars, 1 mm (A and V) or 100 μm (B, C, F-O, R-U, W, and X).
